# Responses of dairy cows following a change in housing system and social group: a living lab experiment

**DOI:** 10.64898/2026.01.30.702528

**Authors:** Catherine Arpin, Marjorie Cellier, Tania Wolfe, Hayda Almeida, Célia Julliot, Marianne Villettaz Robichaud, Abdoulaye Baniré Diallo, Elsa Vasseur

## Abstract

To investigate how the disturbances associated with a relocation to a bedded-pack barn, such as a housing system change, a milking system change and a social regrouping, impacts the behavior of lactating dairy cows, 38 cows from a total of 9 tie-stall or free-stall commercial farms were moved to a newly built bedded-pack barn on an enrollment basis, with a social regrouping occurring after 2 weeks. Scan sampling of video data was done to assess behavior expression in the pen, and live data was collected to assess milking reactivity and animal handling procedures. Results indicate that the cows adapted quickly to the relocation to the new housing system as there were no changes in the locations in the pen, the body positions or the behaviors of cows in time between arrival and regrouping. The social regrouping had a bigger impact with a decrease in 16% of the observed time spent lying and an increase of 9.7% of the observed time spent feeding. Cows also adapted quickly to the milking procedures with a rapid decrease in the occurrence of negative social interactions between cows at the parlor, and in needing less human-animal manipulations and less time to be brought to the parlor. The housing system of origin had a slight effect on behaviors with cows from tie-stalls spending 1.7 times more of the observed time lying than free-stall cows, and free-stall cows spending 1.6 times more of the observed time feeding than tie-stall cows. This study provides a better understanding of how dairy cows respond to disturbances and is encouraging for producers that need to make changes to their current housing system as cows were shown to be quickly adaptable to the challenges presented to them.

**Summary:** Dairy cows from cubicle systems were shown to adapt quickly after a relocation to a bedded pack barn, the first use of a milking parlor, and a social regrouping. This was supported by limited changes observed in their behaviors after the disturbances, and observed deviations were temporary and short-lived. Animal handling procedures also observed a quick improvement in time with the trips to the milking parlor needing 2x less time and 3.5x fewer physical contacts from handlers after 5 days. These results are encouraging to producers needing to make changes to their barns.

## 1. Introduction

Tie-stall housing systems remain the most common housing system type used in Canada with 72% of dairy farms tethering their cows to individual stalls (AAFC, 2024b). This housing system does protect cows from adverse environmental conditions and consequently has been associated with a lower prevalence of lameness (Jewell et al., 2019) than other systems. However, research has demonstrated several health and welfare concerns related to the use of tie-stalls such as higher incidences of knee, hock and neck lesions (Beaver et al., 2021; Mattiello et al., 2009; Regula et al., 2004), less lying time (Beaver et al., 2021; Enriquez-Hidalgo, Teixeira, et al., 2018; Haley et al., 2000), and a higher prevalence of abnormal behaviors such as bar biting, tongue rolling, and leaning against equipment (Beaver et al., 2021; Corazzin et al., 2010; Krohn, 1994).

In order to evolve according to new scientific knowledge and negative public opinion of tie-stalls (Erjavec & Klopčič, 2022; Robbins et al., 2019), there have been unprecedented modifications done to the Dairy Code of Practice, prohibiting the continuous tethering of dairy cows as of April 2027 (NFACC, 2023). This will leave tie-stall farmers in Canada adapting their current system in one of two ways: either modifying their tie-stall barns to allow daily untethered time (such as outdoor access) or adapting their facilities to a less restrictive housing system such as free-stall or bedded-pack barns. This change in the Code will therefore inevitably lead to an increase in the number of cows being relocated to renovated or newly built housing systems and milking systems, as these types of barns often operate using a milking parlor or a robot.

The transition period that occurs after a housing change is of interest from a welfare standpoint as it introduces many new stressors that necessitate behavioral adjustments and could possibly have negative impacts on the health and welfare of the animals. The transition of cows from tie-stall farms to a free-stall barn has been shown to temporarily reduce lying time and milk production (Broucek et al., 2017; Pavlenko et al., 2018, 2023), demonstrating that it could be a stressful period for the animals if not anticipated or mitigated. Understanding how long animals are affected behaviorally by this transition and how long it takes for the cows to get used to the new procedures in place (for example the use of a milking parlor) are of interest in order to develop scientific knowledge to support farmers in transitioning their cattle and to better understand what it entails.

There are not many studies that looked at the effect of a transition from one housing system to another. Most studies between housing systems are comparative, meaning they compare the behavior and other variables between cows housed in different housing systems, which does not take into account the transition and habituation time (Fregonesi & Leaver, 2001; Krohn, 1994; Krohn et al., 1992; Krohn & Munksgaard, 1993; Munksgaard & Simonsen, 1995). From the few studies that moved cows from one barn to the next, they were almost always moving cows from a tie-stall barn to a free-stall barn (Broucek et al., 2017; Pavlenko et al., 2018, 2023). To our knowledge, there has not been a study that brought cows to a bedded-pack farm, which is an emerging type of housing system for dairy cows. Additionally, this study differs by moving cows from both tie-stall and free-stall barns to a bedded-pack system. This project was conceptualized following a living lab approach (Veeckman et al., 2013), meaning that procedures were carefully designed with farmers and animal caretakers in order to represent as much as possible a more realistic scenario that would be observed in a real commercial farm, which makes the conclusions more applicable to real life. Additionally, it tested out a gentler integration to the new barn by adding small groups of cows at a time to mitigate possible stressors caused by the transition. This study also considers not only cows’ behaviors in their pens, but also at the milking parlor, as this is a new environment and procedure for all the cows enrolled in the study.

Additionally, this study looked at the effect of a social regrouping on the behavior of the cows, as regroupings represent a common practice in dairy production. Social regroupings have been shown to affect the behavior of dairy cows by increasing agonistic interactions (Raussi et al., 2005; Schirmann et al., 2011; von Keyserlingk et al., 2008), and decreasing lying times (von Keyserlingk et al., 2008) and feeding times (Schirmann et al., 2011). Furthermore, dairy heifers have been shown to still exhibit increased agonistic interactions after multiple regroupings, never fully getting used to them (Raussi et al., 2005). To investigate the effect of a social regrouping on dairy cows and compare this effect to the one caused by the move to the new barn, this study will also create a regrouping after arrival.

All in all, this study aims to understand how the transition towards a new barn and a new milking system, and a social regrouping are experienced by lactating dairy cows and has two main objectives: (1) Examine how a transition to a new housing system, a new milking system, and a regrouping affects the cows’ behaviors in time, and (2) Assess how previous housing system influences the behavior of cows and their reactivity to milking.

We expect to see changes in the time-budget of the animals immediately after transition (Pavlenko et al., 2018, 2023) such as an initial decrease in feeding and lying and an initial increase in locomotion and exploration, which should stabilize within the first 10 days (Broucek et al., 2017; Pavlenko et al., 2018, 2023). We also expect more agonistic social behaviors right after transition and right after the regrouping as the social hierarchy is established (Raussi et al., 2005; Schirmann et al., 2011). As for milking-related observations, we expect higher milking reactivity from tie-stall-originating cows and higher difficulty with bringing them to the milking parlor than free-stall-originating cows since they are less used to walking and/or responding to humans (Aigueperse & Vasseur, 2021). Finally, we expect a decrease in milking reactivity-related behaviors and difficulty of trips to the parlor within the first weeks as the cows adapt to the new routine.

Overall, this study provides insights into how dairy cows respond to housing transitions and the role previous housing system plays in their adaptation. It increases understanding of how cows react to a new environment and helps support farmers in knowing what to expect during a transition. Finally, it brings forth factors that could affect individual cows’ ability to adapt to a transition, which is in identifying individuals that might be more sensitive to change.

## 2. Material & Methods

### 2.1 Ethical Statement

All procedures and the use of animals were approved by the Animal Care Committee of McGill University and affiliated hospitals and research institutes (Protocol #MCGL-10059).

### 2.2 Animal and Experimental Design

The study was conducted in the newly constructed dairy barn for vocational training of Joyceville Institution (ON), comprised of a straw bedded pack and a parallel milking parlor (accommodating up to 16 cows at a time, 8 on each side), and ran for a total of two months (October 3^rd^ to November 29^th^ 2024). Five pens of 139.3m^2^ with a maximum capacity of 12 cows were used in the project. The cows were fed a total mixed ration (TMR) once a day, with feed pushing twice a day, and had *ad libitum* access to water. They were milked twice a day, once at 6am and once at 4pm.

A total of 38 healthy and non-lame lactating Holstein cows originating from 9 different commercial farms participated in this study. Four of these farms of origin were tie-stall barns, and five were modified free-stall barns, with a resulting total of 13 cows from tie-stalls and 25 from modified free-stalls. None of these cows had experience with milking parlors as the cows in tie-stalls were milked at their stalls, and cows in modified free-stalls were milked either in tie-stalls, in headlocks, or by a robot.

The cows were brought in a total of 7 groups of 4 to 6 cows from their farms of origin to the bedded-pack barn. These 7 groups (here referenced as groups A, B, C, D, E, F & G) were added to the barn (and consequently the trial) on an enrollment basis, meaning that new groups of cows arrived on different dates, being October 3^rd^ (groups A & B), October 10^th^ (groups C & D), October 16^th^ (group E), October 25^th^ (group G) and October 28^th^ 2024 (group F) (Figure 1).

**Figure 1.**
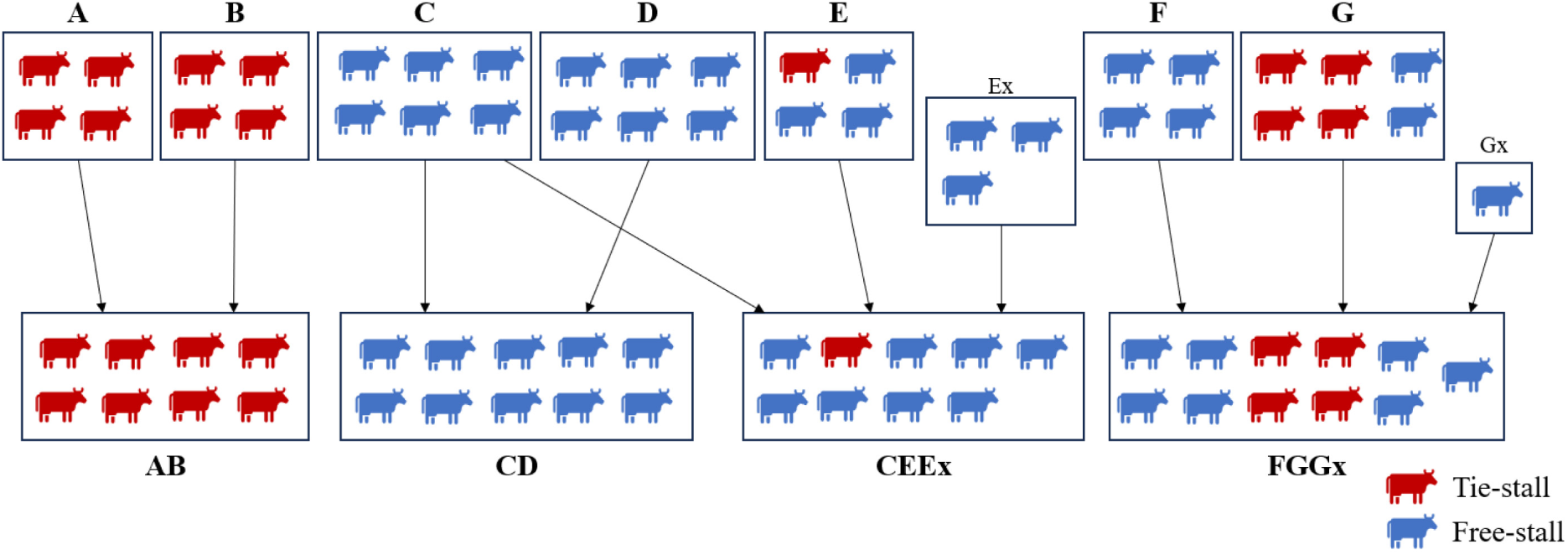
Depiction of the composition of initial groups (A, B, C, D, E, F, G) and the regrouped groups (AB, CD, CEEx, FGGx). Cows from tie-stall barns are shown in red and cows from free-stall barns are shown in blue. Ex and Gx cows were not enrolled in the study upon their arrival, but were added to groups during the regrouping to even the numbers.

For biosecurity reasons, these original groups were kept separate in their pens and were also kept separate during milking for approximately 10 days (between 8-14 days after arrival), limiting interactions between cows from different groups. After around 10 days of separation, groups were combined into bigger groups to form the regrouping. After all regroupings (October 14^th^, October 24^th^ & November 7^th^), there were 4 big groups formed of 8 to 11 cows (here referenced as groups AB, CD, CEEx & FGGx). The exact dates were different for each group as they were introduced on an enrollment basis, but the general timeline depicted in Figure 2 remained the same between groups. *Ex* and *Gx* cows were not initially enrolled in the project, thus were added to the barn separately, kept in separate pens, and were only used to add cows to the regrouped groups (Groups CEEx and FGGx, respectively) (Figure 1). After regrouping, the new groups of cows were studied for an additional 3 weeks, for a total of approximately 5 weeks in the study. Data collection was therefore divided into 3 periods: after arrival (days 1-14), after regrouping (days 15-18), and after stabilization (days 19-38). Stabilization was deemed reached on day 19 (4 days after regrouping) as literature has suggested that cows adapt to a new barn in 10 days (Broucek et al., 2017; Pavlenko et al., 2018, 2023) and a regrouping in 2 days (Raussi et al., 2005; Schirmann et al., 2011; von Keyserlingk et al., 2008).

**Figure 2.**
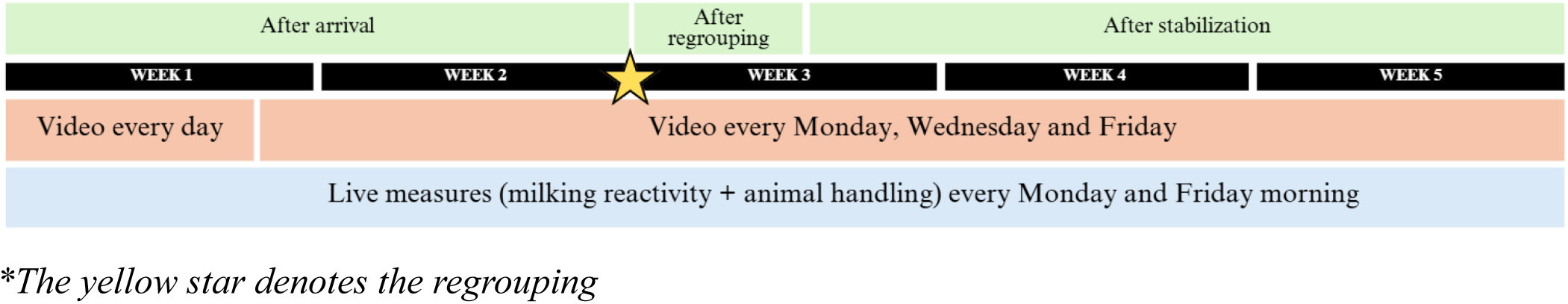
General timeline of the study and data collection for each group. The three periods of study are depicted in green, namely after arrival (days 1-14), after regrouping (days 15-18) & after stabilization (days 19-38). The yellow star demonstrates when the regrouping was held for the group, which was between 8 and 14 days after arrival, depending on the group. The exact dates are different for every group, but the schedule followed these guidelines for each group.

Data collection consisted of video recordings of the pens and live measures taken during milking (see section 2.4 below). Metadata was also collected 3 times per day (once in the morning, once in the afternoon, and once in the evening) for the temperature, the humidity, the wind speed, and the general weather. Temperatures ranged from -1 and 21°C and humidity ranged between 11 and 100%.

### 2.3 Living lab approach

This project was conducted following a living lab approach (Veeckman et al., 2013), meaning that decisions were made closely with farm staff to ensure that the procedures were as manageable as possible and to make the results practical and immediately applicable to commercial farms, going beyond fundamental research.

Additionally, the new farm is a vocational training farm, which involves continuous enrollment and training of individuals with different levels of experience with dairy cows, most having never interacted with them. This implies that procedures were done by many different people with different levels of experience and comfort, and interactions with the animals were not explicitly standardized, instead following a more realistic adoption of practice. Animal handling procedures were adapted over the course of data collection, with animal handling and welfare training offered to the staff and trainees about halfway through data collection, on November 1^st^, with both theoretical and hands-on sessions on how to handle the cows in a safe, controlled and gentle way.

### 2.4 Data collection

#### 2.4.1 Video recording data

Four AXIS-P3227-PVE (AXIS Communications, Inc., ON, Canada) cameras were installed in the barn at a height of approximately 3.5 meters with overlapping views covering the 5 pens. The cameras were wired through a switch (TP-Link model TL-SG1016PE, TP-Link Canada Inc., ON, Canada) to a laptop situated in a secure box in the barn. Recordings were scheduled every day for the first 5 days after the arrival of a group, and 3 times per week until the end of data collection (Monday, Wednesday and Friday; Figure 2). Using the AXIS Camera Station Pro software (Version 6, AXIS Communications, Inc., ON, Canada), cameras were setup to automatically record 3 times per recording day: once in the morning (7:00-9:00), once in the afternoon (11:30-13:30) and once in the evening (16:30-19:30). These times were used to capture the behavior of cows after the morning and evening milkings, and once in the afternoon when there were no humans in the barn, which were assessed to be the most representative of the daily time budget of the cows. Once all recordings were secured, the footage was blurred using an in-house developed algorithm (Araújo et al., 2025) to protect the identity of the workers.

#### 2.4.2 Video annotation

To annotate the recorded videos, scan sampling of the cows’ behavior was done on BORIS software (version 9.3.2) by two observers. All observers completed standardized training, and annotation began only after each observer achieved intra- and inter-observer reliability scores of at least 85% against gold standard. Reliability was checked every 100 videos. Scan sampling was done at 10-minute intervals (following Mitlöhner et al., 2001) for 3h/day, for the full 2 hours in the afternoon slot and 1 hour in the evening slot, starting 1 minute after the cows had access to their whole pen when coming back from the evening milking. Because of the layout of the barn, cows from the two middle pens were confined to their bedded area when returned from milking until all cows were back in their pens. The annotation period started when these cows were given access to their whole pens, and for the other groups, when the gate was closed behind them. The evening timeslot was restricted to only 1 hour to have a total of 3 hours of scan sampling per day, which is similar to what was observed in the literature (Pavlenko et al., 2018, 2023). The morning timeslot was excluded from analysis as the milking end times were quite variable between days, and cows were restricted to the feeding alley at times when the bedded packs were cleaned.

During these scans, the location in the barn (bedded pack or feeding alley), the body position (lying or standing) and the behavior (resting, feeding, locomotion, social, grooming or exploration) of the focal cow were identified based on the ethogram below (Table 4). Annotations were done on all videos included in the timeline of each group (from day 1 to day 38) and were done on a subsample of 30 cows, which were randomly selected from each group according to housing type (4 from A, 4 from B, 5 from C, 4 from D, 4 from E, the Ex cow, 4 from F & 4 from G, resulting in 8 cows from AB, 7 cows from CD, 7 cows from CEEx, and 8 cows from FG) with a resulting 13 cows from tie-stalls and 17 from free-stalls.

Cows were identified on the recordings using pictures taken of all cows upon arrival with a GoPro® HERO11 Black camera (GoPro Inc., San Mateo, CA, USA), one from the front, one from the back, and one from each side.

#### 2.4.3 Live data

##### 2.4.3.1 Milking reactivity

Milking reactivity data was taken 2 times per week during the morning milking for all groups enrolled in the study at the time. Groups were milked one by one, making sure to keep the groups separate. Two observers were used; one on either side of the parlor to simultaneously observe the behavior of cows on both sides, starting when all the cows from the group were aligned in the parlor to when they were released after milking. Behavioral observations were done by noting which of the 10 chosen behaviors were done by each individual (yes or no), including stepping, stomping, kicking, swaying, being reactive to the milking attachment, ruminating, defecating, urinating, vocalizing, and positive and negative social contact (Table 1). The ethogram was created based on previous studies assessing milking reactivity (Brahmi et al., 2024; Jacobs & Siegford, 2012; Marçal-Pedroza et al., 2020; Mincu et al., 2021; Morales-Pineyrua et al., 2023; Sutherland et al., 2012). Observers practiced this method together before the start of the data collection to ensure uniformity.

**Table 1.** Behaviors observed during milking reactivity assessments.

| Behavior | Definition |
| --- | --- |
| Step | The cow lifts a hind leg no higher than 20 cm. |
| Stomp | The cow lifts a hind leg higher than 20 cm. |
| Kick | Backward kick of hind leg (elevating the hind hoof above the hock line and directing it laterally towards the stockperson). |
| Sway | The cow pendulates her weight from one hind leg to the other. |
| Reactive | The cow is reactive to the machine being attached, observed by an increase in movements during the attachment (ex. more steps and kicks). |
| Ruminating | The cow is eating partially digested feed and is either regurgitating or chewing. |
| Defecating | The cow lifts tailhead before eliminating feces from the body. |
| Urinating | The cow arches back and lifts tail to expel urine from the body. |
| Vocalizing | The cow stretches her head forward or upwards with mouth open to produce a call/sound). |
| Social behavior | Positive: The cow makes positive physical contact with neighboring cow (ex. allogrooming, cuddling). |
|  | Negative: The cow makes negative physical contact with neighboring cow (ex. headbutting, pushing). |

*Adapted from Brahmi et al., 2024; Jacobs & Siegford, 2012; Marçal-Pedroza et al., 2020; Mincu et al., 2021; Morales-Pineyrúa et al., 2023 & Sutherland et al., 2012.*

##### 2.4.3.2 Animal handling procedures

There were also multiple variables that were measured relating to the animal handling before, during, and after milking (adapted from procedures by Aigueperse et al., 2023 and Muszik, 2025). This data was taken 2 times per week at the group level during the morning milking for all groups enrolled in the study at the time. For each group, the time needed to bring the cows to the parlor (starts when the first cow takes a step out of the pen and ends when all cows of the group are aligned properly in the parlor), the time needed to milk them (starts when all cows of the group are aligned properly in the parlor and ends when all cows in the group are released from the parlor), and the time needed to bring the cows back to their pen (starts when all cows are released from the parlor and ends when cows are back in their pen and the gate is closed behind them) were taken with a stopwatch. During the trip into the parlor, the types of encouragement needed to bring the cows were observed. These interactions were divided into contact and non-contact interactions (defined in Table 2) and the amount of each were counted. When counting the number of interactions, only the bouts of interactions were considered, meaning that there needed to be a small pause or change of type of interaction to count another one. For example, a member of the staff continuously vocally encouraging the cows counted as one interaction, but if the person stopped and started again, this counted as another interaction. Similarly, if a worker did two types of interactions (ex. pushed a cow and then nudged her leg), this would count as two interactions.

**Table 2.** Definitions of contact and non-contact interactions done by handlers to bring the cows to the parlor.

| Type of interaction | Definition |
| --- | --- |
| Contact | The handler encourages the cow to move forwards by physically manipulating the cow using hand(s), body or object(s). This includes touching the legs from below in the milking parlor, nudging legs, pushing the cows... |
| Non-contact | The handler encourages the cows to move forward by making a noise such as clapping, whistling, tongue clicking, speaking etc... or waving of the hand(s), arm(s) etc... |

*Adapted from Aigueperse et al., 2023 and Muszik, 2025.*

### 2.5 Statistical analysis

#### 2.5.1 Data handling

##### 2.5.1.1 Video data

The raw measures for each ethogram component (ie., location in the barn, body position and behavior) extracted from the annotated videos consisted of the frequency of the observed ethogram component for each cow in each video, which was then transformed into ratios. For each cow, ratios were calculated by dividing the frequency of every ethogram component by the total number of annotated scans in the video to give a ratio of the time spent doing each component. For example, if a cow was lying for 6 of the 12 scans, we considered that the cow lied down for 50% of the video. Ratios for afternoon and evening timeslots were calculated separately and not summed together, as not every day had both timeslots due to data loss.

##### 2.5.1.2 Live data

For the milking reactivity analysis, no transformations were done and data was kept as binary, where 1 being the cow did the behavior during milking and 0 being the cow never did the behavior during milking. For the animal handling procedures, the number of contact and non-contact interactions were kept as numbers, but the timed variables (ex. time needed to get the cows to the parlor) were transformed into meters per second, dividing the distance covered between the pen and the parlor by the time to standardize the variable across pens. Analysis of variables before and after the regrouping was done separately without direct comparison between them as they were group measures, and groups changed after regrouping.

#### 2.5.2 Data analysis

All descriptive statistics were done on Excel (version 2504), and all statistical analysis were performed on R® (version 4.4.3 (2025-02-28 ucrt) -- “Trophy Case” (250 Northern Ave, Boston, MA 02210)) using glmmTMB, lme4, lmerTest, nlme and emmeans packages for model analysis. Assumptions of normality (when needed) were graphically assessed using Q-Q plots and histograms, and when not met, appropriate transformations were applied (Table 3). For repeated measures, the best-fitting covariance structure was assessed, with AR(1) model providing the best fit based on BIC values. To accommodate the AR(1) covariance structure, the data were nested to ensure a single observation per time point. Consequently, we acknowledge that the degrees of freedom are higher than standard models due to this specific structural requirement. For post-hoc pairwise comparisons, tukey or scheffe corrections were applied. Significance was tested using anovas and was declared at P ≤ 0.05, with tendencies falling between 0.05 and 0.1. Results are reported using generated emmeans.

**Table 3.** Data transformation to reach normality of residuals.

| **Ethogram category** | **Analysis** | **Transformation** |
| --- | --- | --- |
| **Ethogram component** |  |  |
| Live data |  |  |
| Speed (m/s) |  |  |
| Speed to the parlor | N/A | none |
| Speed back to the pen | N/A | none |
| Frequency of human-animal interactions |  |  |
| Contact interactions | N/A | sqrt |
| Non-contact interactions | N/A | log |

| Video data (% of scans for each component) |  |  |
| --- | --- | --- |
| Location in the barn | In time after arrival | arcsin sqrt |
| Bedded pack, feeding alley or border | Between periods | logit |
|  | Between housing systems | arcsin sqrt |
| Body position | In time after arrival | logit |
| Lying, standing or moving | Between periods | logit |
|  | Between housing systems | none |
| Behavior | In time after arrival | sqrt |
| Resting/inactive, feeding, locomotion, social, exploration or grooming | Between periods | logit |
|  | Between housing systems | arcsin sqrt |

#### 2.5.3 Live data

To assess the effect of time and farm type of origin on the live data variables (i.e. milking reactivity behaviors, speed in, speed out, contact and non-contact interactions), linear mixed models were used.

##### 2.5.3.1 Milking reactivity

To account for the binary nature of the data (yes/no), a generalized linear mixed model with the binomial family was used, with fixed effects including the Farm type of origin and Day, and the random effect of Cow. The model was used for every reactive behavior measured (step, stomp, kick, sway, reactive to the milking attachment, rumination, defecation, urination, vocalization, positive social contact, and negative social contact) and was as follows:

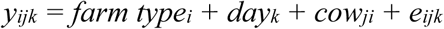

Because of the use of a generalized model, the normality of residuals was not needed, but an overdispersion test was done on the residuals for each milking reactivity behavior.

##### 2.5.3.2 Animal handling procedures

The model was similar for the animal handling portion with fixed effects including the Farm type of origin and Day, but the random effect being the Group. The following model was applied to Speed In (to the parlor), Speed Out (back to the pen), and the number of contact and non-contact human interactions needed to bring the cows to the parlor:

#### 2.5.4 Video data

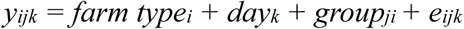

There were three objectives with this portion of analysis: (1) Testing if there were changes in the expression of ethogram components (ex. lying, feeding, socializing) in time after arrival, (2) Testing if there were differences in the expression of ethogram components between the 3 periods (after arrival, after regrouping, and after stabilization), and (3) Comparing the expression of ethogram components between cows originating from tie-stall barns and cows originating from free-stall barns. Three similar linear mixed models were used to investigate the objectives across the 3 ethogram categories: Location, Body Position and Behavior, which were included as fixed effects. All models included random effects accounting for Timeslot and Cow, though the exact nesting structure varied by model. An autoregressive correlation structure of order 1 (Ar1) was used to account for the repeated measures.

##### 2.5.4.1 Differences in time after arrival

To assess the variation in the expression of ethogram components in time, Day, Ethogram category, and their interaction were included as fixed effects. A random intercept of Timeslot nested within Group and Cow was included to account for repeated measures and individual variation. The following model was done for each Ethogram category:

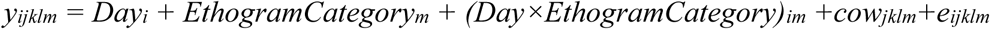

Where cow_jklm_ is the random intercept and e_ijklm_ are the residuals modeled with Ar(1) autocorrelation over Day.

**Table 4.**
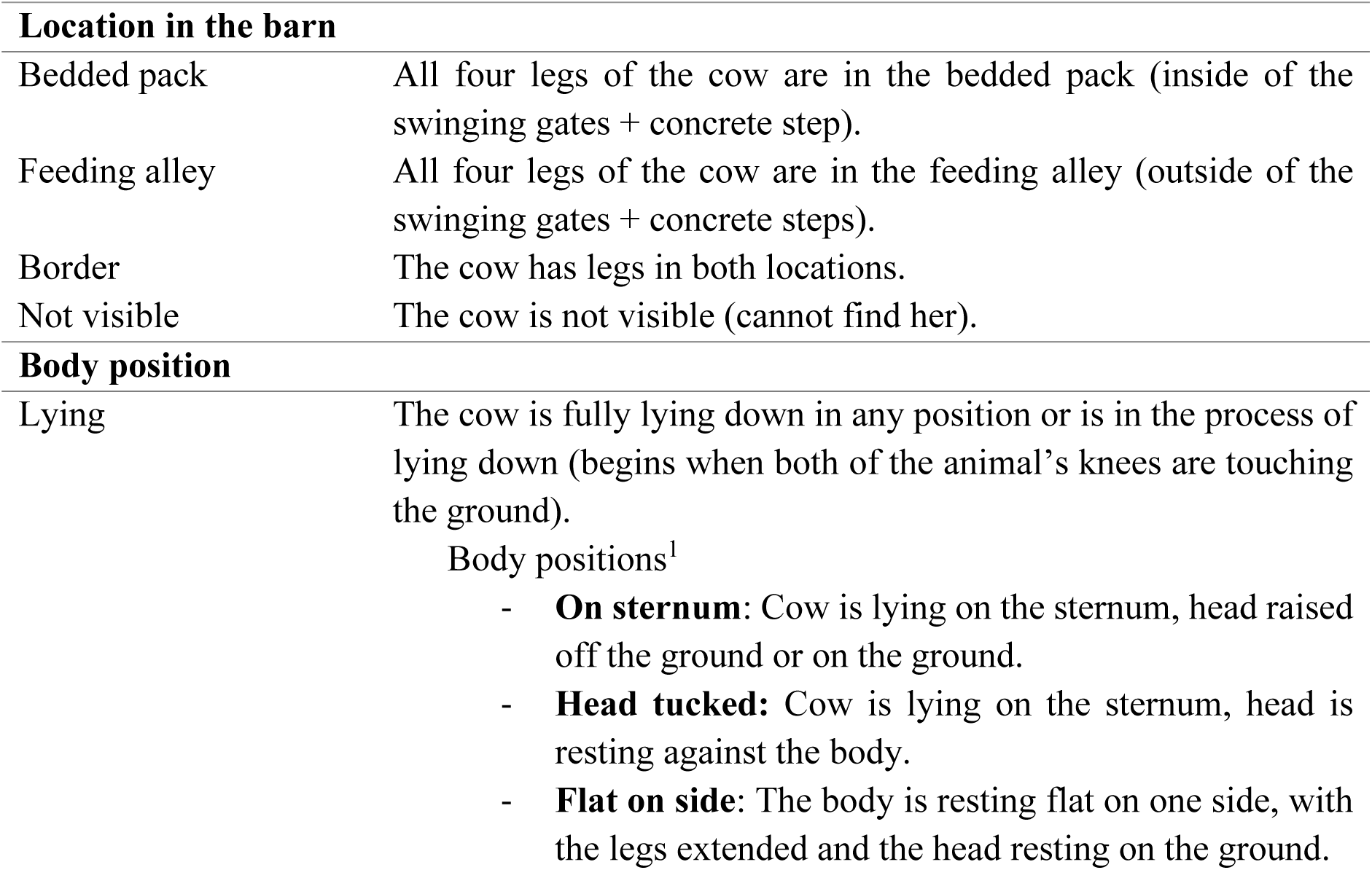
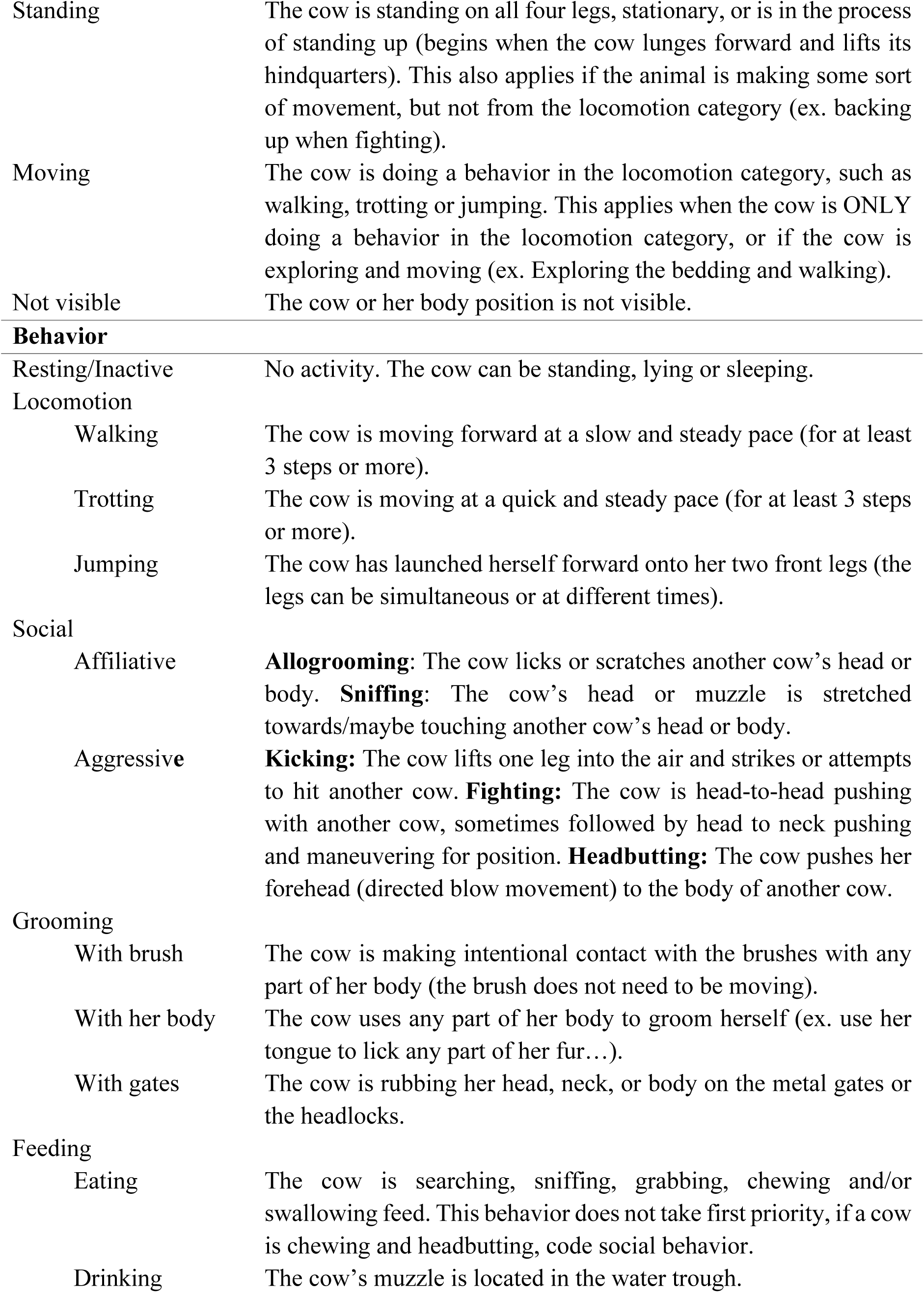
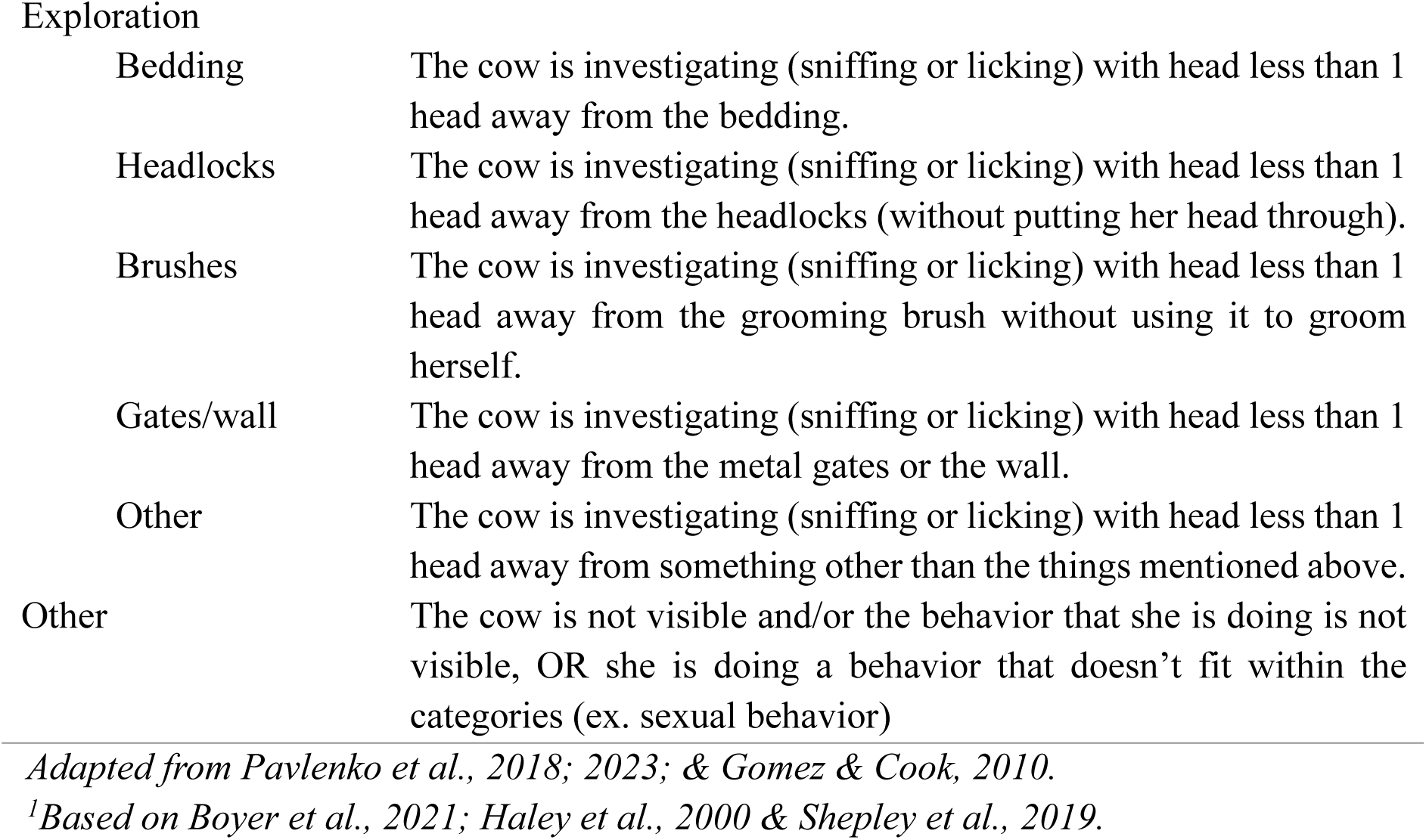
Ethogram used to annotate videos. Location, Body position & Behavior were separate categories, meaning that for every scan, one object from each category was coded.

| **Location in the barn** |  |
| --- | --- |
| Bedded pack | All four legs of the cow are in the bedded pack (inside of the swinging gates + concrete step). |
| Feeding alley | All four legs of the cow are in the feeding alley (outside of the swinging gates + concrete steps). |
| Border | The cow has legs in both locations. |
| Not visible | The cow is not visible (cannot find her). |
| **Body position** |  |
| Lying | The cow is fully lying down in any position or is in the process of lying down (begins when both of the animal's knees are touching the ground).<br>Body positions<sup>1</sup><br>- **On sternum:** Cow is lying on the sternum, head raised off the ground or on the ground.<br>- **Head tucked:** Cow is lying on the sternum, head is resting against the body.<br>- **Flat on side:** The body is resting flat on one side, with the legs extended and the head resting on the ground. |
| Standing | The cow is standing on all four legs, stationary, or is in the process of standing up (begins when the cow lunges forward and lifts its hindquarters). This also applies if the animal is making some sort of movement, but not from the locomotion category (ex. backing up when fighting). |
| Moving | The cow is doing a behavior in the locomotion category, such as walking, trotting or jumping. This applies when the cow is ONLY doing a behavior in the locomotion category, or if the cow is exploring and moving (ex. Exploring the bedding and walking). |
| Not visible | The cow or her body position is not visible. |
| Behavior |  |
| Resting/Inactive | No activity. The cow can be standing, lying or sleeping. |
| Locomotion |  |
| Walking | The cow is moving forward at a slow and steady pace (for at least 3 steps or more). |
| Trotting | The cow is moving at a quick and steady pace (for at least 3 steps or more). |
| Jumping | The cow has launched herself forward onto her two front legs (the legs can be simultaneous or at different times). |
| Social |  |
| Affiliative | **Allogrooming:** The cow licks or scratches another cow's head or body. **Sniffing:** The cow's head or muzzle is stretched towards/maybe touching another cow's head or body. |
| Aggressive | **Kicking:** The cow lifts one leg into the air and strikes or attempts to hit another cow. **Fighting:** The cow is head-to-head pushing with another cow, sometimes followed by head to neck pushing and maneuvering for position. **Headbutting:** The cow pushes her forehead (directed blow movement) to the body of another cow. |
| Grooming |  |
| With brush | The cow is making intentional contact with the brushes with any part of her body (the brush does not need to be moving). |
| With her body | The cow uses any part of her body to groom herself (ex. use her tongue to lick any part of her fur...). |
| With gates | The cow is rubbing her head, neck, or body on the metal gates or the headlocks. |
| Feeding |  |
| Eating | The cow is searching, sniffing, grabbing, chewing and/or swallowing feed. This behavior does not take first priority, if a cow is chewing and headbutting, code social behavior. |
| Drinking | The cow's muzzle is located in the water trough. |
| Exploration |  |
| Bedding | The cow is investigating (sniffing or licking) with head less than 1 head away from the bedding. |
| Headlocks | The cow is investigating (sniffing or licking) with head less than 1 head away from the headlocks (without putting her head through). |
| Brushes | The cow is investigating (sniffing or licking) with head less than 1 head away from the grooming brush without using it to groom herself. |
| Gates/wall | The cow is investigating (sniffing or licking) with head less than 1 head away from the metal gates or the wall. |
| Other | The cow is investigating (sniffing or licking) with head less than 1 head away from something other than the things mentioned above. |
| Other | The cow is not visible and/or the behavior that she is doing is not visible, OR she is doing a behavior that doesn't fit within the categories (ex. sexual behavior) |

*Adapted from Pavlenko et al., 2018; 2023; & Gomez & Cook, 2010.*
*<sup>1</sup>Based on Boyer et al., 2021; Haley et al., 2000 & Shepley et al., 2019.*

##### 2.5.4.2 Differences between periods

To assess the differences in the expression of ethogram components between periods, the Period, the Ethogram category, and their interaction were included as fixed effects. A random intercept of Period nested within Day, Timeslot and Cow was included to account for repeated measures and individual variation. The model was as follows:

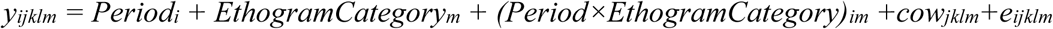

Where cow_jklm_ is the random intercept and e_ijklm_ are the residuals modeled with Ar(1) autocorrelation over Period.

##### 2.5.4.3 Differences between housing systems of origin

For the comparison of ethogram components between farm types of origin, this was done more a posteriori, as this was not accounted for in the original experimental design. As this was a living lab experiment and the origin of cows was not something that could be controlled, the fact that cows came from either tie-stall or free-stall barn was noticed during data collection but was not originally planned. However, the distinction between cows in either farm type proved to be of interest, and the statistical analysis was done using the model described below. Housing system of origin (HS), ethogram category, and their interaction were included as fixed effects. A random intercept of HS nested within Day, Timeslot and Cow was included to account for repeated measures and individual variation. The model was as follows:

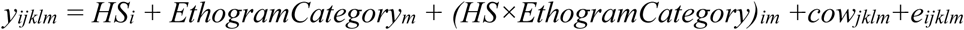

Where cow_jklm_ is the random intercept and e_ijklm_ are the residuals modeled with Ar(1) autocorrelation across Day.

## 3. Results

### 3.1 Changes in time

#### 3.1.1 Behaviors after arrival

In the 14 days between arrival and regrouping, cows had a similar percentage of scans in the bedded pack (54.2±4.2%) and in the feeding alley (45.5±4.2%; p=0.309). There were no significant differences or trends observed between days, but a high variability with the percentage of scans in the bedded pack falling between 36 and 66% and the percentage of scans in the feeding alley falling between 31 to 64% (Figure 3). Body position-wise, cows spent significantly more of the observed time standing (57.5±4.4%), than lying (37.1±4.2%; p=0.004). There were observed twice as often lying on day 4 than day 1 (21.9±4.8%; 51.7±6.6%; p=0.009), but no other significant differences or trends were found within lying or the other body positions. There was also a high variability in lying and standing body positions with cows spending between 22 and 52% of the observed time lying and between 42 and 77% of the observed time standing. Behavior wise, cows spent significantly more of the observed time resting (40.8±2.6%) than feeding (26.7±2.1%; p<0.001) and spent little of the observed time exploring (2.8±0.7%), grooming (0.1±0.1%), socializing (0.2±0.2%) and moving (0.6±0.3%). There was significantly less feeding observed on day 4 than on day 3 (14.5±3.0%; 34.2±4.0%; p=0.035), but no other observations or trends observed between days. Behaviors were found to be more stable in time with less variation between days.

**Figure 3.**
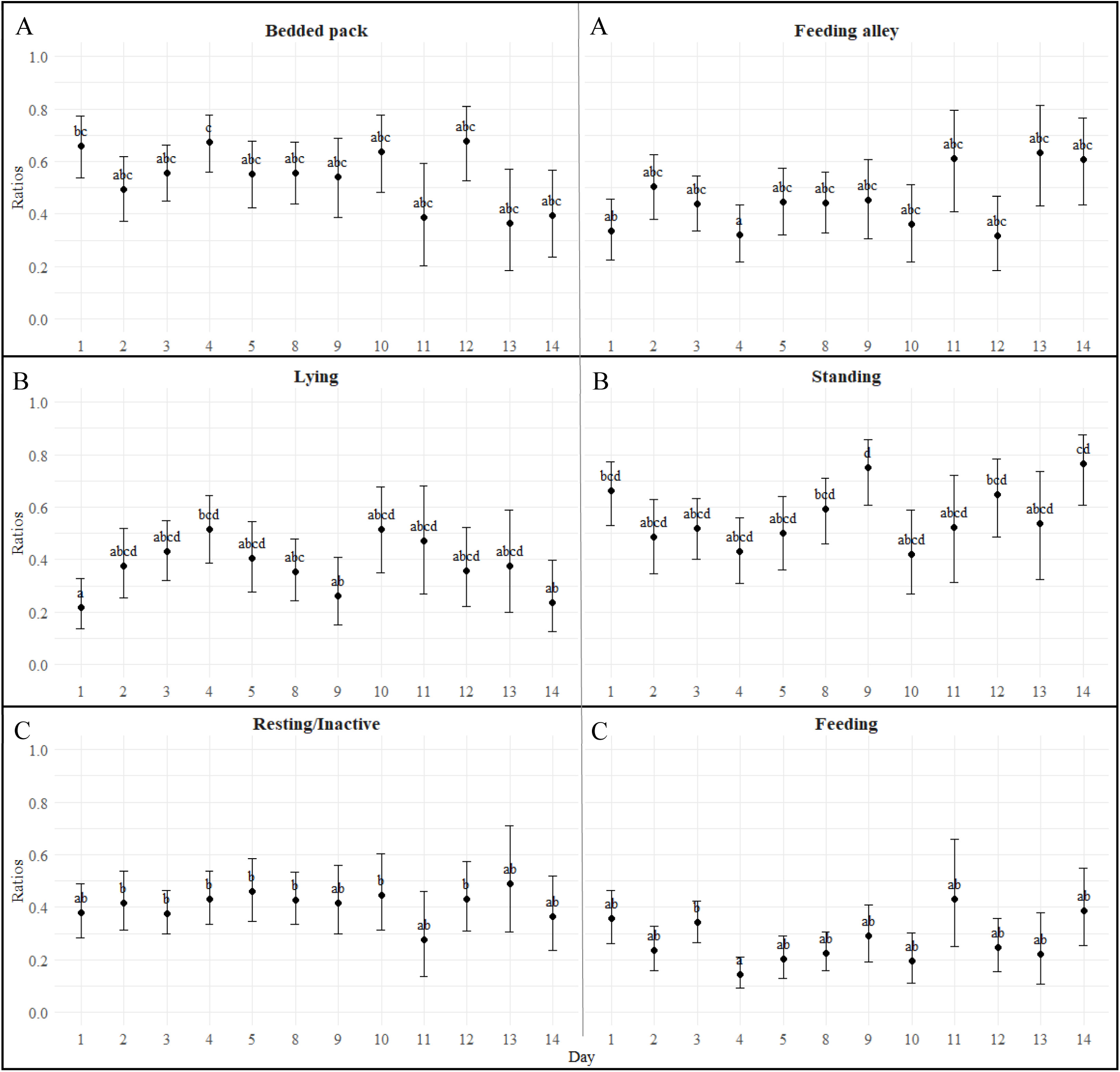
– Evolution of the ratios of scans spent doing each ethogram component by day for the 14 days between arrival and regrouping (after arrival period). Letters above the points designate statistically significant differences between days (days without a common letter are statistically different: Tukey post-hoc test for pairwise comparisons, p < 0.05). Graphs with the same big letter have been compared statistically between them, meaning the little letters apply to them both.

#### 3.1.2 Behaviors between periods

In this section, how cows allocated their time was compared between periods (after arrival, after regrouping & after stabilization), as they were found to be stable between days in each separate period (Figure S1). Regrouping was found to have had an impact on many behaviors. Compared to the period after arrival, location-wise, the percentage of scans in the bedded pack were reduced by 8.8% after regrouping (p=0.112, df = 2070, t-ratio = 2.00), which was mirrored by an increase of 7.8% in the feeding alley after regrouping (p=0.174, df = 2070, t-ratio = -1.79) (Table 5). Body positions were also different between the after arrival and after regrouping periods with a decrease of 16.8% in scans spent lying after regrouping (p<0.001, df = 2070, t-ratio = 3.95) and an increase of 16.3% in scans spent standing (p=0.0014, df = 2070, t-ratio = -3.50). Finally, behavior-wise, only resting and feeding behaviors were found with differences between after arrival and after regrouping periods, leaving exploration, grooming, social and locomotion without inter-period variation. The percentage of scans spent resting tended to decrease by 8.2% (p=0.0574, df = 4080, t-ratio = 2.29) and the percentage of scans spent feeding significantly increased by 9.7% after regrouping (p=0.0069, df = 4080, t-ratio = -3.03). In all cases, percentage of scans after arrival and after stabilization were not statistically different.

**Table 5.**
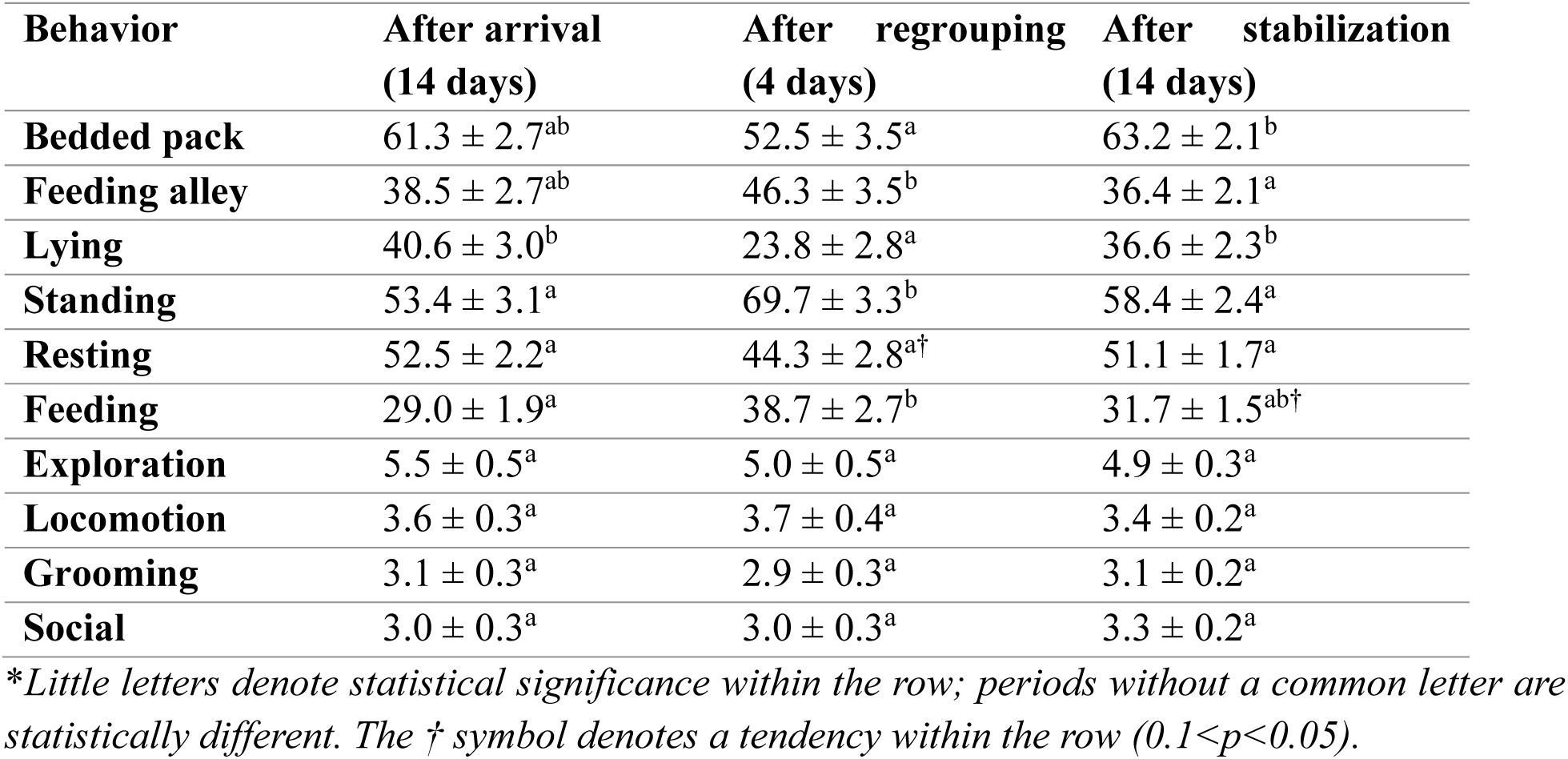
Percentages of scans spent doing each specific ethogram component for each period, after arrival (which lasted 14 days), after regrouping (which encompassed the 4 days after regrouping), and after stabilization (which includes 14 days).

#### 3.1.3 Milking reactivity and animal handling procedures in time

Regarding reactive behaviors done at milking, the only behavior that varied in time was the occurrence of negative social interactions, such as headbutting and fighting. Cows had a significantly higher probability of having negative social interactions during milking on the first week of data collection (59.0 ± 9.5%) than the 4th (22.4 ±7.5%; p=0.025) and 5th (9.8 ±4.8%; p<0.001) weeks, showing that the chances of aggressive behaviors happening at milking decreased in time (Figure 4). However, the probabilities of stepping, stomping, kicking, vocalizing, urinating, defecating, and having positive social contacts stayed significantly stable in time.

**Figure 4.**
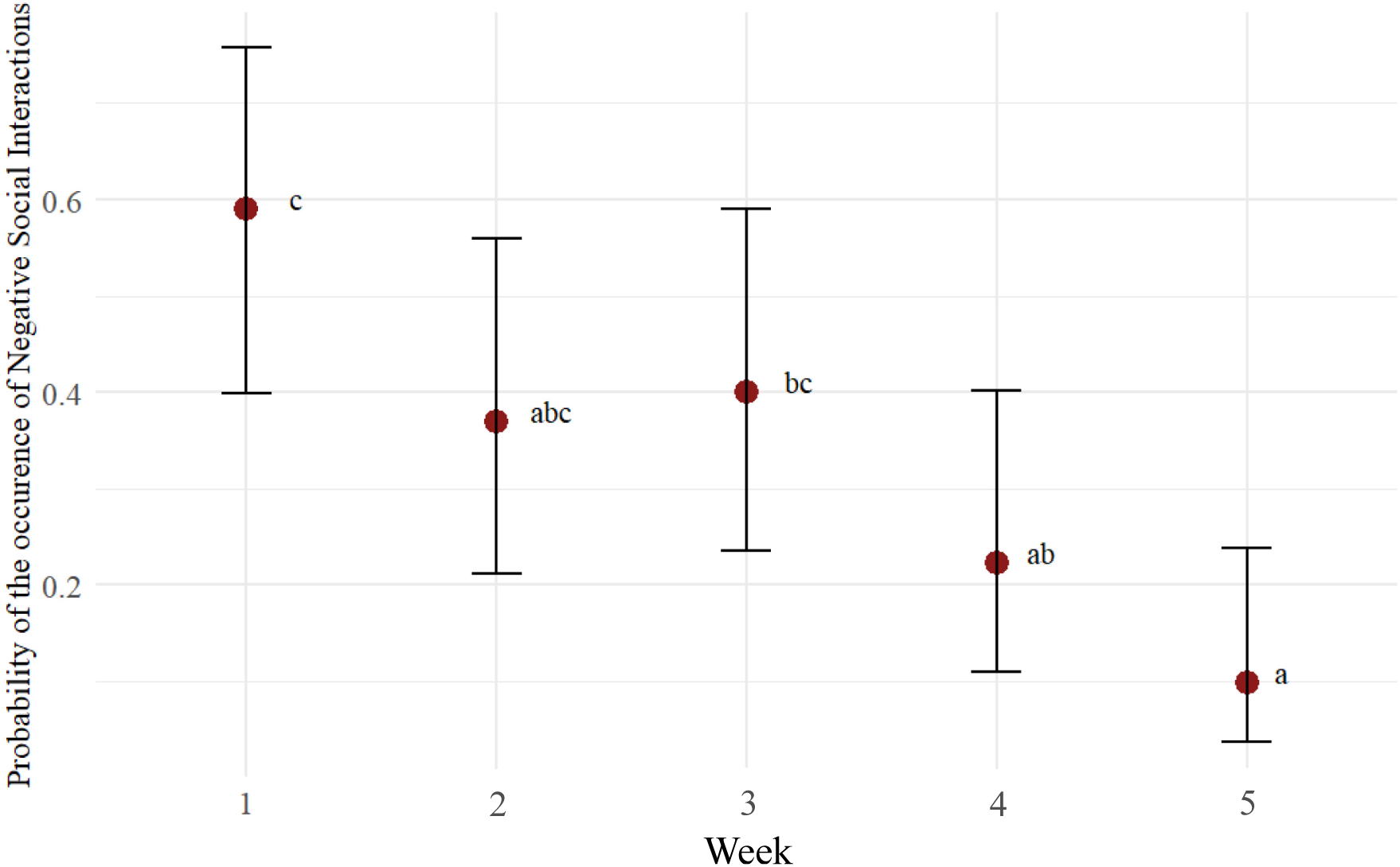
Probability of negative social interactions occurring during milking for each week of the experiment. All groups were observed on slightly different days during the week, within a 4-day window (week 1 = days 2-5; week 2 = 9-12; week 3=16-19; week 4 = 23-26; week 5 = 30-33).

As for the animal handling procedures, cows rapidly got faster at getting to the parlor. They were over 2 times faster on the 5^th^ day (p=0.102, df = 7.81, t-ratio = -2706) and 3 times faster on the 12^th^ day (p=0.0058, df = 7.80, t-ratio = -4.872), compared to the 2^nd^ day. When looking at the human-animal interactions needed to bring the cows to the parlor, they rapidly decreased with over 3.5 times less contact interactions needed on day 5 (p=0.202, df = 11.4, t-value = 2.127) and 7 times less contact interactions on day 12 (p=0.0279, df = 10.6, t-value = 3.382) (Figure 5).

**Figure 5.**
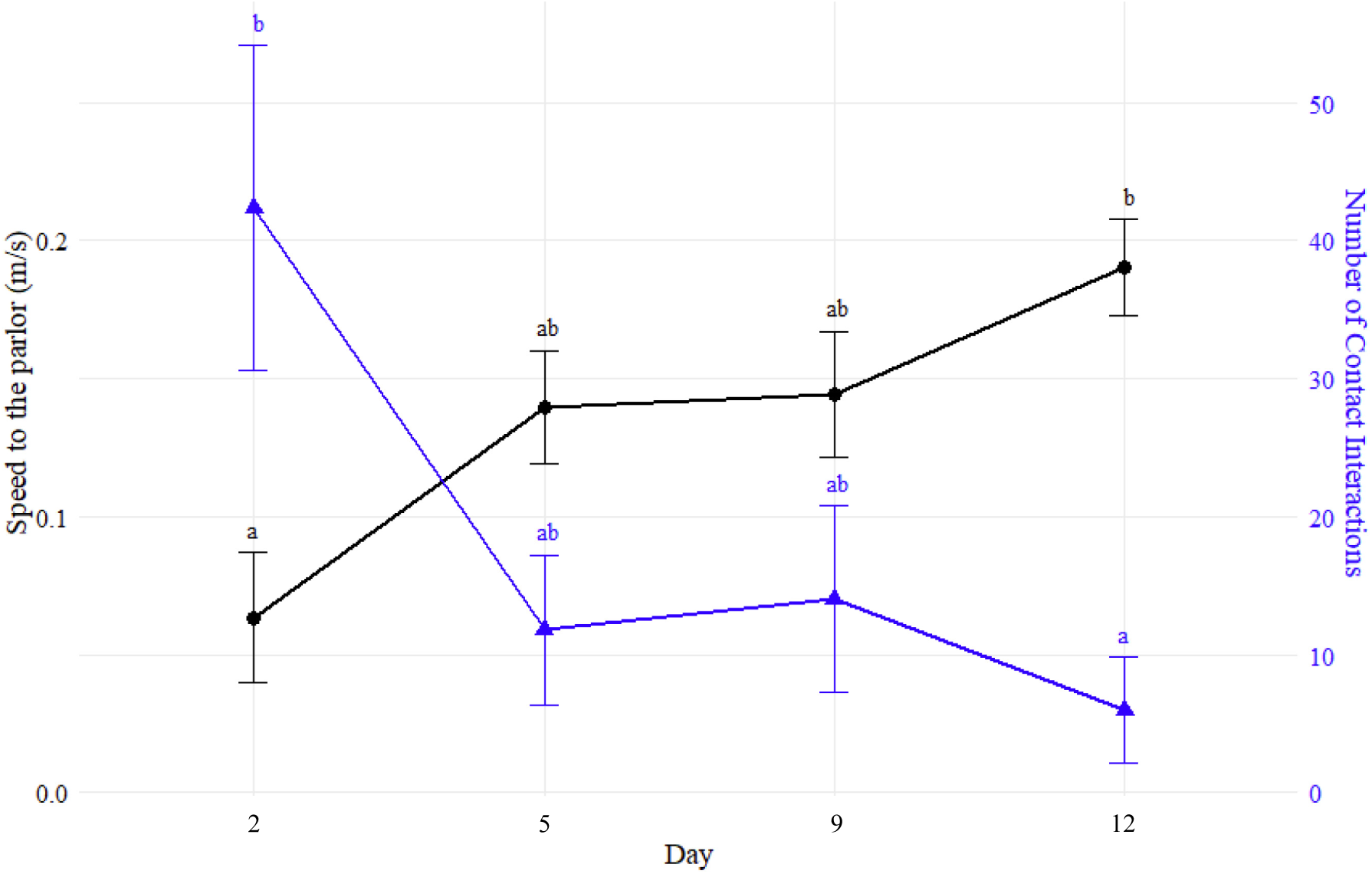
Evolution of the speed of cows (m crossed/time taken) (black circles) and number of contact human-animal interactions (blue triangles) during the trip to the milking parlor in time. Letters above the points designate statistically significant differences between days for that variable (days without a common letter are statistically different: Tukey post-hoc test for pairwise comparisons, p < 0.05). Speed and number of interactions were analyzed separately.

### 3.2 Effect of previous housing system

In this section, cows were identified as having been brought from a tie-stall barn (TS) or a free-stall barn (FS) and their behaviors in the pen and reactivity during milking from the beginning to the end of the trial are compared.

#### 3.2.1 Comparing behaviors of TS and FS cows

Regarding the location of the cows in the barn, FS cows spent the same percentage of the observed time in both locations (p=1), compared to TS cows that spent 2.7 times more of the observed time in the bedded pack than in the feeding alley (p<0.0001). There were also differences in the percentages of scans spent in each body position. FS cows were found standing 2 times more often than lying (p<0.0001), meanwhile TS cows spent 1.3 times more of the observed time lying than standing (p<0.0001). As for the specific behaviors, FS and TS cows spent similar limited amounts of the observed time socializing, exploring, grooming and in locomotion, but were different in their distribution of percentage of scans spent resting and feeding. Similarly to the location, FS cows were found resting and feeding in similar percentage of scans (p=1), compared to TS cows that were found resting 2.5 times more often than feeding (p<0.0001).

There was also quite a variety in how the cows did their behaviors (Table 6). For example, all 3 lying positions were observed with the position lying on the sternum being prioritized and found 15.7 times more than the other positions combined. As for social behaviors, allogrooming, sniffing, fighting and headbutting were all observed in scans with sniffing and fighting being done in the greatest number of scans. Grooming was done with the brushes, the gates and their bodies in similar proportions. Finally, cows explored every aspect of their environment such as the bedding, the headlocks, the brushes, the gates, the water trough (in Other) etc. with the greatest number of scans spent exploring the gates.

**Table 6.** The percentage of scans spent in each subcategory of behaviors for lying, social, grooming and exploratory behaviors. The totals in bold represent the total number of scans annotated for each behavioral category (ex. lying). The percentages were calculated based off that total.

| **Behavioral category** | **Sub behavior** | **TS** | **FS** |
| --- | --- | --- | --- |
| **Lying** | **Total** | **1382** | **1280** |
|  | On sternum | 94.0% | 94.3% |
|  | Head tucked | 5.1% | 5.4% |
|  | Flat on side | 0.9% | 0.3% |
| **Social** | **Total** | **35** | **64** |
|  | Allogrooming | 8.6% | 23.4% |
|  | Sniffing | 34.3% | 15.6% |
|  | Fighting | 37.1% | 43.8% |
|  | Headbutting | 20.0% | 17.2% |
| **Grooming** | **Total** | **27** | **53** |
|  | Brush | 37.0% | 32.1% |
|  | Body | 51.9% | 35.8% |
|  | Gates | 11.1% | 32.1% |
| **Exploration** | **Total** | **155** | **203** |
|  | Bedding | 25.2% | 18.7% |
|  | Headlocks | 12.9% | 22.2% |
|  | Brushes | 1.3% | 1.0% |
|  | Gates | 57.4% | 54.2% |
|  | Other | 3.2% | 3.9% |

#### 3.2.2 Comparing milking reactivity and animal handling procedures of TS and FS cows

There were not many differences between TS and FS cows when comparing the probability of different reactive behaviors at milking being done by the cows. The two behaviors that were significantly different between TS and FS cows were swaying and positive social contacts. TS cows were 1.6 times more likely to sway during milking than FS cows, and 2.1 times more likely to engage in positive social contacts (Table 7). There were no differences in stepping, stomping, kicking, vocalizing, urinating, defecating, and negative social contacts between housing systems.

**Table 7.** Probability of each reactive behavior being done by cows during milking.

| **Reactive behavior** | **TS** | **FS** | **p-value** |
| --- | --- | --- | --- |
| **Stomp** | 42.0 ± 8.7% | 36.5 ± 5.8% | 0.589 |
| **Kick** | 25.6 ± 9.5% | 19.1 ± 5.7% | 0.538 |
| **Sway** | 64.3 ± 7.2% | 40.2 ± 5.2% | 0.01 ** |
| **Reactive** | 24.8 ± 7.9% | 14.5 ± 3.9% | 0.195 |
| **Rumination** | 21.8 ± 5.1% | 23.6 ± 3.9% | 0.777 |
| **Positive Social Contact** | 53.1 ± 6.4% | 25.0 ± 3.7% | 0.0001 *** |
| **Negative Social Contact** | 21.3 ± 5.6% | 31.1 ± 4.8% | 0.194 |

*\*Asterisks denote statistical significance*
*Step, Vocalization, Defecation & Urination were removed on cases of quasi-complete separation, caused by the fact that the probability of vocalization, defecation and urination were too close to 0 and step was too close to 100%.*

As for the animal handling data, housing system of origin had no significant impact on the time needed to bring the cows to the parlor or the number of human-animal interactions of either type needed to get them to the parlor.

## 4. Discussion

Original hypotheses were partially supported by the results described in this paper. There were no observed changes in the expression of behaviors in pen after the arrival to the new barn, and no increase in social behaviors after the social regrouping, contrary to predictions. The trips to the milking parlor did improve quickly in time, as predicted, with cows needing fewer physical interactions from handlers and taking less time to reach the parlor, however, no differences were detected between TS and FS cows in this analysis. The two groups were found to have some differences, primarily in their main activity in their pens (TS cows spending more time resting and FS cows spending more time feeding), and during milking, with TS cows swaying more often than FS cows.

### 4.1 Examine how a transition to a new housing system, a new milking system, and a social regrouping affects the cow’s behaviors in time

When comparing the location in the barn, the body position and the behavior of cows in time between arrival and regrouping, there were no clear trends or variation observed. The only behaviors that had significant differences between days were lying (with an increase between days 1 and 4) and feeding (with a decrease between days 3 and 4). However, in both cases, the general changes between days were quite sporadic and stayed relatively within normal ranges, with lying staying within 22-52% of the observed time (Tucker et al., 2021) and feeding remaining between 14.5-43% of the observed time (Campler et al., 2018, 2019; Fregonesi & Leaver, 2001, 2002; Laínez & Liang, 2004; Munksgaard & Simonsen, 1995; Pereira et al., 2023; Phillips & Schofield, 1994). This, combined with the lack of differences of the other behaviors seems to indicate that the cows in this study adapted quite immediately to the new housing system.

It is difficult to compare these results to the literature since to our knowledge, there haven’t been studies done observing the transition period after a relocation to a bedded pack barn. However, the move from a tie-stall barn to a free-stall barn has been measured in the past (Pavlenko et al., 2018, 2023). In these two papers, large groups of cows were added to the barn at the same time with 400-960 cows in the barn within the first 5 days, compared to our 8 cows on the first day (and week). The metrics used were also slightly different than ours, as they used the percentage of cows doing each behavior at each scan, compared to our individual measures, but the sampling times were similar to our 3 hours per day and 10 min interval scans. In both studies, changes were observed in time after arrival. In the 2018 study, authors observed no changes in lying and standing between weeks, similarly to our study, but in the 2023 study, they observed an immediate increase of 20% in standing and a 20% decrease in lying during the first week. Both studies observed different levels of increase in walking after arrival (1.5 and 10%) and either a decrease of 15% in feeding or no differences in time. Slight differences were also found in time for grooming, exploration and aggression. Since we observed no such differences or trends in time, this seems to suggest that our methodology of introducing small groups at a time, compared to the two described studies, and keeping them separate in order to limit regroupings was effective in making this transition period as smooth and least stressful possible. Additionally, it shows that even in a less controlled environment with frequent interruptions, employees in training and quick changeover of personnel, which is more reflective of actual adoption of practice, cows were still able to adapt quickly.

Moreover, the distribution of time spent between behaviors established was similar to the ones described in other studies, with lying and feeding times being within normal ranges. However, it is important to mention that our sampling consisted of scan sampling at 10-min intervals for 3 hours a day, not a full 24-hour time-budget, meaning that percentages might not be fully comparable. Average lying times in lactating dairy cows have been reported as being between 8 and 13 hours, or 33-54% a day (Tucker et al., 2021) and approximately 39% of the day in a bedded pack barn (Endres & Barberg, 2007), which are very similar to our 37%. Feeding times in our study (27%) were also within the range described in other studies of 15-40% (Campler et al., 2018, 2019; Fregonesi & Leaver, 2001, 2002; Laínez & Liang, 2004; Munksgaard & Simonsen, 1995; Pereira et al., 2023; Phillips & Schofield, 1994). Even with scan sampling, our percentages falling within range shows that the behaviors of the cows were done in “normal” proportions during the chosen sampling times, and could indicate that cows were in fact adapted to their new housing system.

Compared to the arrival to the new barn, the regrouping seemed to have had more of an impact on the cows with an observed decrease in the percentage of the observed time spent lying of 16.8% and an increase in feeding of 9.7%. These are similar to results observed in other studies measuring the effects of a social regrouping on the behavior of dairy cows (Raussi et al., 2005; von Keyserlingk et al., 2008), which observed a decrease in lying times and no changes in feeding times after regrouping. However, we observed no differences in social behaviors, compared to most regrouping studies that report an increase in social behaviors, especially aggressive behaviors (Raussi et al., 2005; von Keyserlingk et al., 2008). Additionally, our regroupings also brought an increase in density of cows as the groups got bigger (from 4-6 to 8-11 cows per pen), which has been shown to increase competition (Fregonesi et al., 2007), and either decrease (Fregonesi et al., 2007; Jurkovich et al., 2024) or have no effect on lying times (Fregonesi & Leaver, 2002; Mogensen et al., 1997). However, in our study, the increase in density was not accompanied by resource limitations, such as lack of lying space or lack of available headlocks, which was created in most of the studies cited above. To avoid this, the number of cows per pen was kept under the total capacity of 12 cows. The abundance of resources could explain why no changes in social behaviors were observed after the regrouping. Another explanation could be that since our data was collected using scan sampling, behaviors that occupy less of the time-budget of cows, such as social behaviors, are harder to catch in scan sampling (Mitlöhner et al., 2001) and therefore making it more difficult to observe slight trends. Even if changes in behaviors such as feeding and lying were observed, percentages were kept close to the normal range, and all behaviors were done in their proper place. For example, lying was kept to the bedded pack as there were no reports of cows lying in the feeding alley. Additionally, the period after regrouping consisted of only 4 days and percentages were back to pre-regrouping levels in the next period, which shows that the effect was short-lived as there were no significant changes in time detected within periods (Figure S1). Therefore, the social regrouping did have an effect, but temporarily and not to the point of being necessarily detrimental to their welfare.

As for the changes of milking reactivity in time, we noticed a decrease in the probability of negative social interactions between cows occurring at milking in time. This could have been due to the gradual stabilization of the hierarchal structure in time (Boe & Færevik, 2003; Broucek et al., 2017) and cows choosing their order in the parlor. It appears that no other reactive behavior significantly changed in time, showing that cows adapted very quickly to being milked at the parlor. In the future, it would be interesting to look at milk production metrics in time to get a bigger picture of the stress response and effects on milk production, especially since a drop in milk production has been observed after a transition from a tie-stall barn to a free-stall barn (Pavlenko et al., 2018, 2023).

We also noticed that cows got faster at getting to the parlor as they got used to the trip, which has been observed in past papers looking at motivation for outdoor access (Aigueperse et al., 2023). This was easily visually observable as cows started waiting at the gates of their pens when the parlor got ready in the morning, making the process of getting them into the alley and to the parlor easier, faster, and more efficient (personal observation). In time, cows also went by themselves and settled into the parlor without help from the handlers. We also observed that handlers got more comfortable and better at handling the cows, which resulted in less unnecessary force used and a resulting decrease in contact interactions needed. This progression was natural and should be representative of the changes in time observed in a commercial setting, which highlights once again the strength of the living lab approach. Staff members and trainees were also offered handling and welfare training halfway through the course of the study, as several of them were manipulating cows for the first time. This training encouraged the use of non-contact interactions and taught the basis of cows’ blind spots and pressure zones, and discouraged unnecessary physical contact, which could have also helped in reducing the observed numbers.

It was difficult to find studies to compare our results to as there weren’t any that directly looked at the transition from milking at the stall to milking in a parlor. However, there were some studies that looked at the transition from a conventional parlor to an automated parlor. These studies noticed a rapid adaptation to the new milking procedures with elimination and vocalization during milking decreasing back to normal amounts within 24 hours (Jacobs & Siegford, 2012) and a stabilization in step counts after 5 days (Morales-Pineyrua et al., 2023). Because of the study design, we were unable to get the milking reactivity data for the first milking for most of the groups, making subtle changes difficult to catch. Additionally, we didn’t measure the frequency of behaviors, but only if they were done or not by the cows. Therefore, behaviors with very low frequencies such as defecation, urination and vocalization, or very high frequencies such as stepping, were impossible to analyze with our models even if there were potential significant differences. Defecation and urination have been shown to increase with fear and stress (Tonooka et al., 2022), so our very small number of reported eliminatory behaviors during milking is a good sign.

Generally, cows seemed to adapt quickly and rapidly to a relocation to a new housing system, a regrouping and a change in milking system. There were no changes observed in the time-budget or the milking reactivity in time after arrival, and even changes observed after regrouping were settled after 4 days. And these results were found even in a context where there is continuous training of staff with newcomers, a fast trainee turnover, and less standardization in the methods with every worker being different. This type of experiment is very important as it is representative of a real transition in a commercial setting, and these results demonstrate that cows can adapt remarkably quickly to big changes and the transition can be a success if it is done properly (i.e. in small groups, with limited regroupings and with handling training offered).

### 4.2 Assess how previous housing system influences the behavior of cows and their reactivity to milking

As explained in the methods section, the comparison between cows originating from tie-stalls and cows originating from free-stalls was more exploratory as the groups were discovered during data collection and therefore the experimental design did not account for this separation of cows from two groups. Nonetheless, interesting results appeared, such as the fact that TS cows and FS cows differed in their main activity. TS cows spent more of the observed time resting, which meant that they were more often found lying, and therefore more often found in the bedded pack. FS cows, however, spent more of the observed time feeding, and therefore were found more often standing and in the feeding alley. These differences could have been attributed to a multitude of reasons, such as FS cows being used to having to walk to get from the lying area to the feeding area, compared to TS cows that were fed at their stalls and were not used to having to move to another location. Additionally, TS cows had no prior experience standing on a concrete floor (as found in the feeding alley), which is less comfortable than standing in a bedded pack (Shepley, Lensink, & Vasseur, 2020), which could explain why TS cows spent less time in the feeding alley than FS cows, that had experienced it before. And in both TS and FS cows, 16% of the observed time spent standing was in the bedded pack, where the cows could have been exploring, grooming or socializing, compared to standing in the feeding alley which was often linked with feeding. There were no differences observed between TS and FS cows when it came to exploration, grooming, social and locomotive behaviors. This is probably because feeding and resting occupy over 90% of cattle’s time-budget (Dickson et al., 2022; Hedlund & Rolls, 1977), which makes it easier to detect differences, especially with scan sampling and with a smaller sample size (Mitlöhner et al., 2001). In order to have more accurate reporting of those behaviors, either continuous sampling or a higher sample size would be required (Mitlöhner et al., 2001).

Both groups were observed interacting with every aspect of their environment such as exploring the bedding, the headlocks, the brushes and the gates, and using the brushes and the gates for grooming. They were also observed doing a variety of social behaviors such as sniffing, allogrooming, headbutting and fighting. Observing such a large diversity of behaviors executed by cows is a sign of positive welfare as it shows that the environment is fulfilling many behavioral needs (L. Miller et al., 2020). The new environment offers many opportunities for exploration, grooming and social behaviors and cows are taking advantage of them.

As for reactive behaviors during milking, TS cows were observed swaying during more milkings than FS cows. This could be a sign of higher discomfort to milking (Medrano-Galarza et al., 2012; Mincu et al., 2021; Raoult et al., 2021) or walking (Shepley et al., 2020) compared to FS cows that are used to having to walk to another area to get milked. It could also be a sign of impatience/restlessness (Cooper et al., 2007) since they are not used to having to wait for all cows to be done milking before getting released, compared to FS cows. There were also more milkings with positive social contacts occurring between TS cows than FS cows, potentially due to needing more interactions to reinforce social bonds and reduce social tension (Boissy et al., 2007) than FS cows. Other than those behaviors, there were no differences between TS and FS cows when it came to milking reactivity and animal handling procedures, showing that cows will adapt quickly to the use of a milking parlor no matter the farm type of origin.

## 5. Conclusion

In conclusion, no matter their housing system of origin, dairy cows adapt quickly to a relocation to a new barn, the use of a new milking system, and a social regrouping. Our methodology was shown to be effective in making this transition as smooth as possible, even among variability inherited from the living lab approach. Overall, this study is encouraging to producers that will need to make big changes to their current housing system following the changing legislation. In the future, collecting behavioral data before and after regrouping for comparation purposes and establishing 24h time-budgets would be beneficial to get a better idea of the effects of these disruptions on the time-budget of dairy cows.

## CRediT Authorship Contribution statement

**Catherine Arpin**: Conceptualization, Data Curation, Formal Analysis, Investigation, Methodology, Visualization, Writing – Original Draft, Writing – Review & Editing. **Marjorie Cellier**: Conceptualization, Formal Analysis, Investigation, Methodology, Supervision, Visualization, Writing – Review & Editing. **Tania Wolfe**: Conceptualization, Formal Analysis, Investigation, Methodology, Supervision, Visualization, Writing – Review & Editing. **Hayda Almeida:** Conceptualization, Methodology. **Célia Julliot**: Conceptualization, Investigation, Methodology. **Marianne Villettaz Robichaud**: Conceptualization, Writing – Review & Editing. **Abdoulaye B. Diallo**: Funding Acquisition, Project Administration, Supervision, Writing – Review & Editing. **Elsa Vasseur**: Conceptualization, Funding Acquisition, Methodology, Project Administration, Supervision, Writing – Review & Editing.

