## Supplemental table for "Responses of dairy cows following a change in housing system and social group: a living lab experiment"

Supplemental Material

**Figure S1** – Evolution of the ratio of scans spent doing each ethogram component in time for the 2 other periods: after regrouping and after stabilization. Day represents day after regrouping. Little letters denote statistical significance between points in the same category (divided by dashed line).

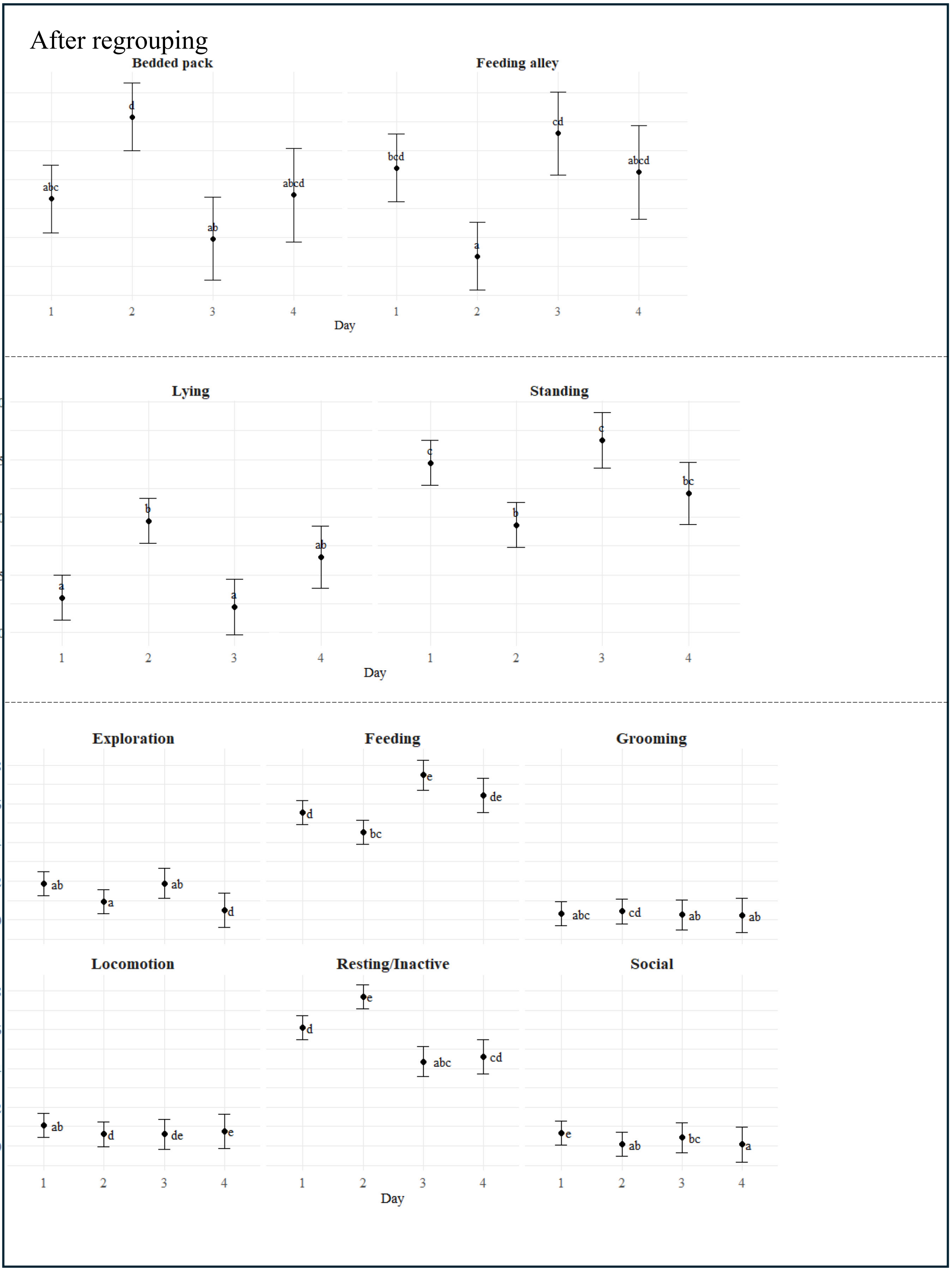

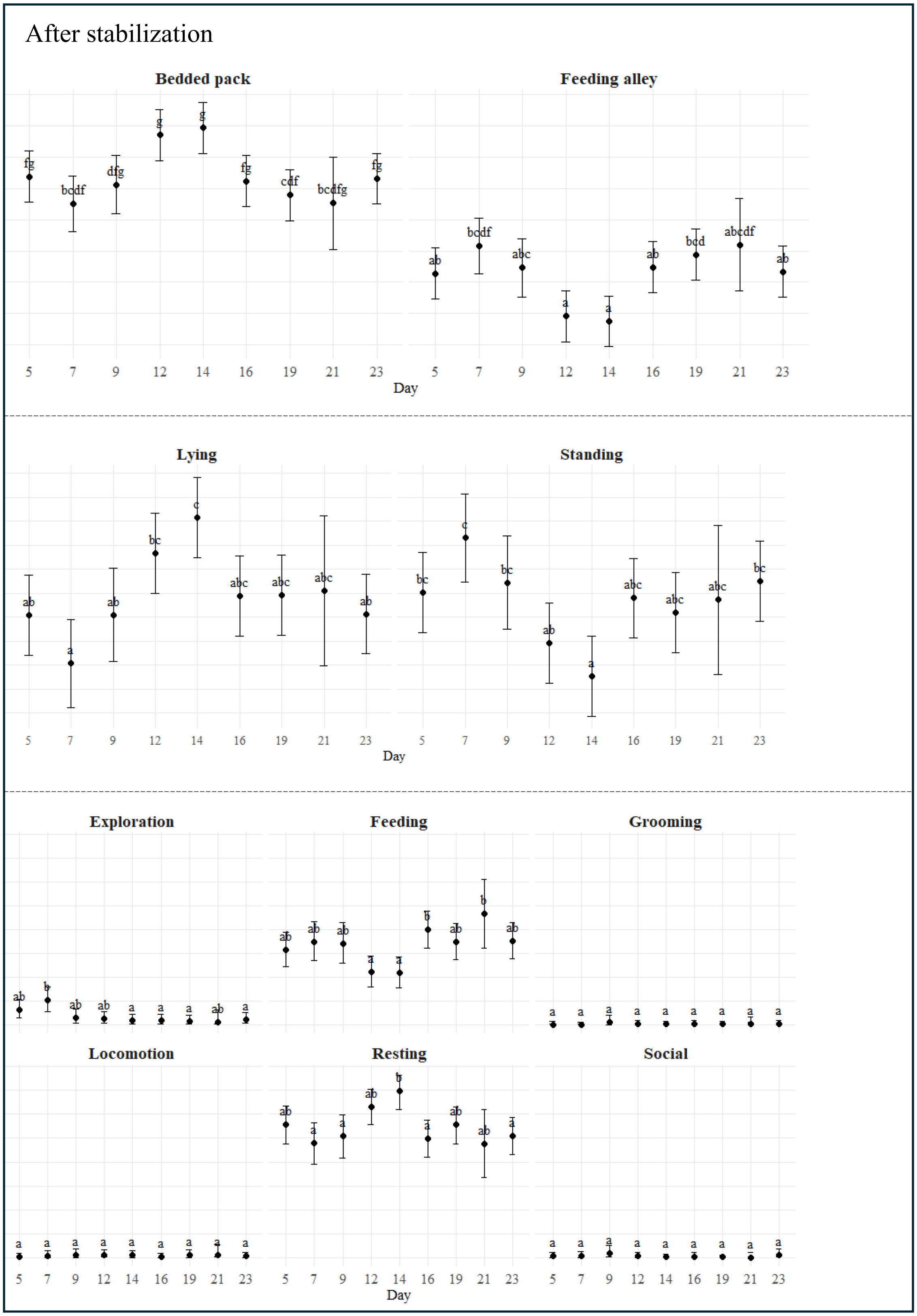

After stabilization
Bedded pack
Feeding alley
Ratios (arcsin sqrt)
Day
Lying
Standing
Ratios (logit)
Day
Exploration
Feeding
Grooming
Locomotion
Resting
Social
Ratios (arcsin sqrt
Day
